# Identification and Characterization of Compounds that Improve Plant Photosynthesis and Growth under Light Stress Conditions

**DOI:** 10.1101/2024.04.25.591172

**Authors:** Yuchen Qu, Kazuma Sakoda, Yu Wakabayashi, Masatoshi Nakajima, Tadao Asami, Ichiro Terashima, Wataru Yamori

## Abstract

In order to satisfy the food and fuel demands of a growing population, global food production needs to increase by more than 50% before 2050. However, various environmental stresses in the natural environment inhibit plant growth and result in reduced yields. This is primarily caused by decreases in photosynthetic capacity. Thus, there is an urgent need to develop new strategies to improve agricultural productivity and ensure food security. In this study, a novel chemical-screening system with 96 well plates and leaf disks of tobacco was used to identify several anthraquinone derivatives that could relieve high light stress from plants. Treatments with these chemicals induced greater photosynthetic capacity after high light stress conditions for 20–72 hours (h) in tobacco and better plant growth after exposure to light stress for 96 hours in Arabidopsis and lettuces. The photoprotective effect of anthraquinone derivatives is closely related to chemical induced oxidation of PSI. Furthermore, there were no negative effects on plant growth in chemically treated plants under non-stressful conditions. Taken together, this study shows that anthraquinone derivatives can confer high light stress tolerance in plants, resulting in improved plant photosynthesis and growth in environments with light stress.

## Introduction

The world’s population may reach 10 billion by 2050, but around 10% currently suffer from food shortages (UN, 2019; FAOSTAT, 2019). Furthermore, natural disasters, such as droughts, floods, and extreme temperatures caused by climate changes, are now becoming a threat to agricultural production (Rosenzweig et al., 2001), and 91% of farmland is impacted by some type of stress, which will eventually result in decreases in grain yields (Minhas et al., 2017). Decreases in plant productivity are primarily caused by reduced photosynthetic ability because of stress damage (Yamori et al., 2016; Sharma et al., 2020). It is well known that plants are sensitive to changes in their environment, and fluctuations in natural conditions may result in stress and hinder the growth and yield of plants by up to 50% (Yamori et al., 2014; Yamori, 2016; Yamori and Shikanai, 2016; Kang et al., 2017; Mosa et al., 2017; Boonchai et al., 2018). Such conditions include drought (water stress), excessive watering (water logging), extreme temperatures (cold, frost, and heat), salinity, and mineral toxicity. Nevertheless, increased light intensity (light stress) induced by climate changes plays an role which cannot be neglected (Hussain et al., 2021). Therefore, improving photosynthesis under stressful conditions is especially important to ensure agricultural productivity and food safety in the near future (Qu et al., 2023).

As an energy source, light is critical for photosynthesis. Plants absorb light at photosystem II (PSII) and I (PSI), giving impetus to the flow of electrons through the electron transport chain, and they eventually turn light energy into ATP and NADPH. ATP and NADPH are consumed in the Calvin cycle, where CO_2_ is reduced and generally assimilated in the form of organic compounds. However, plants frequently get exposed to high light intensities and are photodamaged, resulting in them being unable to utilize all the captured light energy for photosynthesis (Demmig-Adams and Adams, 1992; Long et al., 1994; Takahashi and Murata, 2005). When electron consumption in the CO_2_ assimilation process is no longer sufficient to cope with electron generation, excessive energy starts to accumulate in chloroplasts, resulting in light stress (Demmig-Adams and Adams, 1992; Long et al., 1994; Takahashi and Murata, 2005). Usually, this results in the generation of reactive oxygen species (ROS), which are induced by direct reductions of O_2_ (Demmig-Adams and Adams, 1992; Telfer, 2014), or PSI receptor side over-reduction, leading to a decrease in PSI oxidization capacity due to P700 rapid charge recombination (Endo et al., 1999). In fact, PSII damage from light stress is often observed in field conditions (Ishida et al., 2014), and it is generally considered that PSII is a primary target for photoinhibition as a result of ROS accumulation (Murata et al., 2007). Studies have shown that PSI damage caused by fluctuating light during the day or over-reduction of the electron transport chain in high light conditions is relatively common and can result in decreased photosynthesis, growth, and yields (Ilić et al., 2012; Allahverdiyeva et al., 2015; Yamori et al., 2016; Van Rooijen et al., 2017; Li et al., 2020). One major area of research for improving plant tolerance against light stress is through genetic modifications. There has been much research into the biosynthesis of carotenoids (Davison et al., 2002), flavonols (Emiliani et al. 2013), chloroplastic prenyl lipids (Ksas et al. 2015), and melatonin (Lee and Back 2018) combined with modifications to the xanthophyll cycle (Götz et al. 2002). Recent studies have shown that the introduction of maize *GOLDEN2-LIKE* genes into rice reduced photoinhibition and improved vegetative biomass and grain yields by 30–40% in strong light conditions (Li et al., 2020), and introducing flavodiiron proteins of *Physcomitrium patens* into rice or Arabidopsis reduced photodamage during PSI in high light and fluctuating light conditions (Wada et al., 2018; Basso et al., 2022).

However, there are increasing concerns about the safety of planting and consuming genetically modified plants (Kramkowska et al., 2013). An alternative strategy for improving plant tolerance of light stress could be through chemical biology. Chemical biology is a discipline of biology that addresses biological phenomena by using small bioactive chemicals together with conventional biological techniques (Kitahata and Asami, 2011). In the past two decades, such techniques have been widely explored with animal materials, from the cellular level to high-throughput screening methods for drug discoveries (Giacomotto and Ségalat, 2010; Lessman, 2011). And recently, these techniques have also been used in plant research. Some chemical compounds have been found to be capable of inducing stomatal closure through ABA-dependent or ABA-independent signaling pathways (Kinoshita et al., 2021; Vaidya et al., 2019), while other chemical compounds show commercial potential as candidate compounds for increasing the shelf life of cut flowers in the floral industry (Toh et al., 2018). It has also been found that some chemicals can be utilized in the modification of essential proteins in the regulation of cell division and polarity, which improves stomatal development (Sakai et al., 2017; Bian et al., 2020) and increases seedling biomass in Arabidopsis (Ziadi et al., 2017). Furthermore, treatment with acetate triggers a dynamic metabolic flux conversion from glycolysis into acetate synthesis, which stimulates the jasmonate signaling pathway. This confers drought tolerance and improves the survival of Arabidopsis, maize, wheat, rice, and rapeseed in extreme drought conditions (Kim et al., 2017). However, despite these many breakthroughs, it seems that chemical compounds that regulate plant tolerance against light stress have barely been studied.

In this study, we constructed a novel chemical screening system to search for chemicals that could improve plant photosynthesis and growth under high light conditions. Tobacco leaf disks were used as the major plant material. Screening through chemical libraries was performed, based on photosynthetic imaging techniques. After identifying several anthraquinone derivatives that could relieve high light stress from plants, we further analyzed the underlying mechanisms.

## Results

### Screening a library of 12,000 chemicals

To identify compounds that affect plant tolerance under light stress conditions, we first screened 12,000 chemical compounds acquired from Maybridge using the tobacco leaf disk system. Tobacco leaves have extremely large surface areas with uniformly featured photosynthetic activities, which can enable measuring the effect of the chemicals on a large scale. The entire screening process was divided into first and second screening sections. First screening was performed with one replication for each chemical in order to achieve high throughput, and any chemical with a suspiciously positive effect would be further tested in the second screening. For the second screening, each chemical was tested with three or more replications.

After two rounds of screening, 33 candidates that induced higher Fv/Fm or Y(II) at the initial or stable state of photosynthetic induction (actinic light is 200 µmol m^-2^ s^-1^ followed by 700 µmol m^-2^ s^-1^) in leaf disks after high light stress treatment were identified out of the 12,000 chemicals. Among these candidates, nitrogen dioxide (-NO_2_) residue was found in 27 chemicals and cyanide (-C≡N) residue in six chemicals, making them dangerous or environmentally toxic (Dash *et al*., 2009; Elsayed, 1994). Two anthraquinone derivatives (A4N, A18Ch) were considered to be generally safe and were selected for further study since they showed high potential in practical applications (Figure 1C). In further analyses, we also used 10 structural analogs (A14N, A48N, A1N4C, A1N, A1458N, A1Ch, A14Ch, A14Ch23C, A1Ch2C, A4Ch1C) closely related to A4N and A18Ch (Figure 1C).

**Figure 1.**
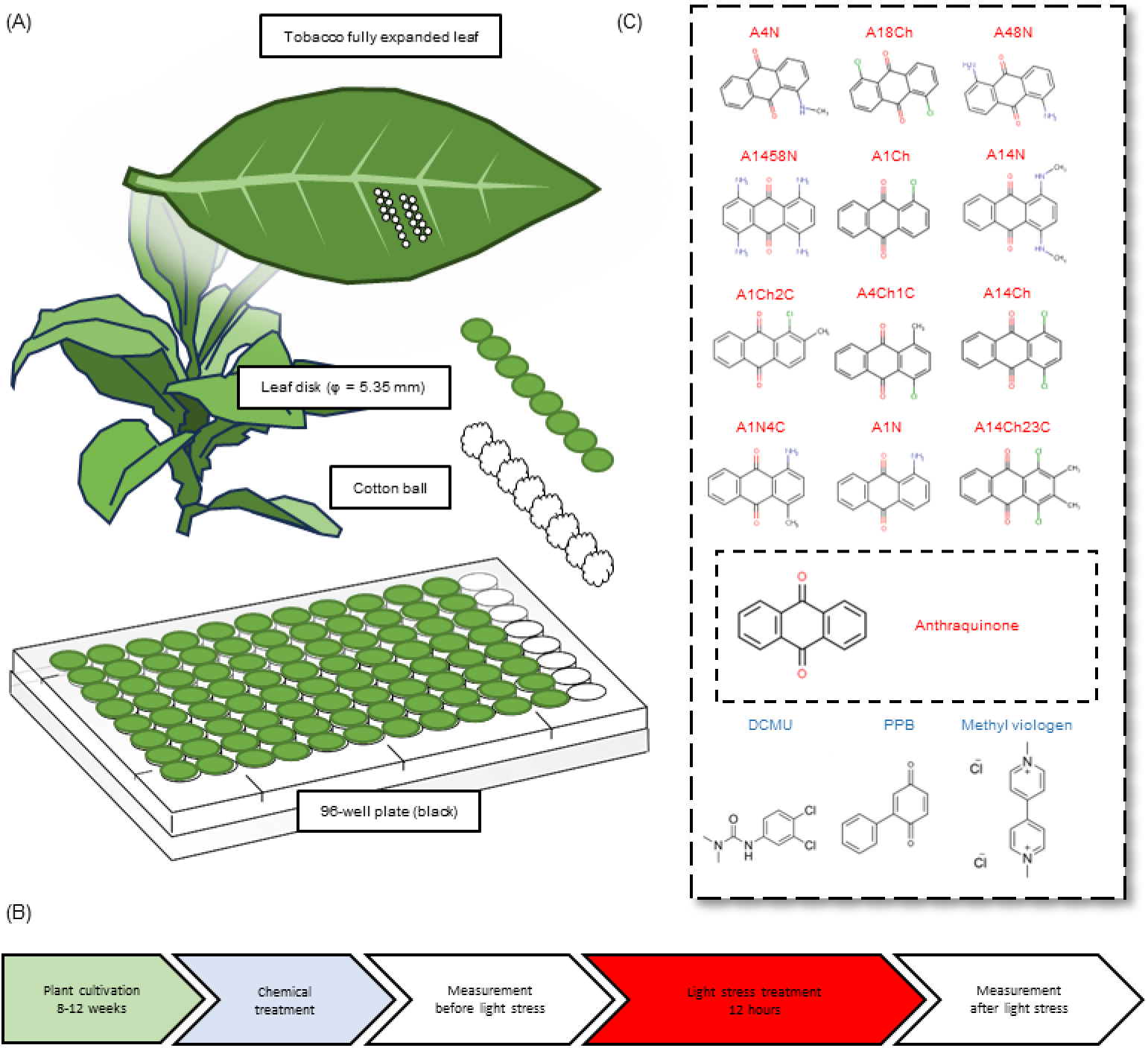
Constructing a screening system using tobacco leaf disks. **A**. Leaf disks and 96 well plates were used as chemical screening tools in this study. **B**. The process for performing the leaf disk chemical screening experiment. **C.** Structural information of candidate chemicals that showed protective effects against light stress, together with three inhibitors/electron acceptors, which served as positive/negative controls.

### Chlorophyll fluorescence of leaf disks before and after light stress treatment

The leaf disks were treated with 12 chemicals, and chlorophyll fluorescence was measured before and after high light stress at 700 μmol photons m^−2^ s^−1^ (Figure 2). A18Ch, A1Ch and A14Ch significantly decreased Fv/Fm even without light stress compared with the control group (Figure S1A). A similar effect was observed in the DCMU and MV treatment groups (Figure S1A). After light stress treatment, the A18Ch, A1Ch and A14Ch treatment groups together with the DCMU treatment group showed significantly lower Fv/Fm compared with the control group (Figure S1B). The MV treatment group showed completely undetectable Fv/Fm (Figure S1B). Other chemical treatment groups showed no significant differences in Fv/Fm compared with the control group before and after light stress (Figure S1A and S1B).

**Figure 2.**
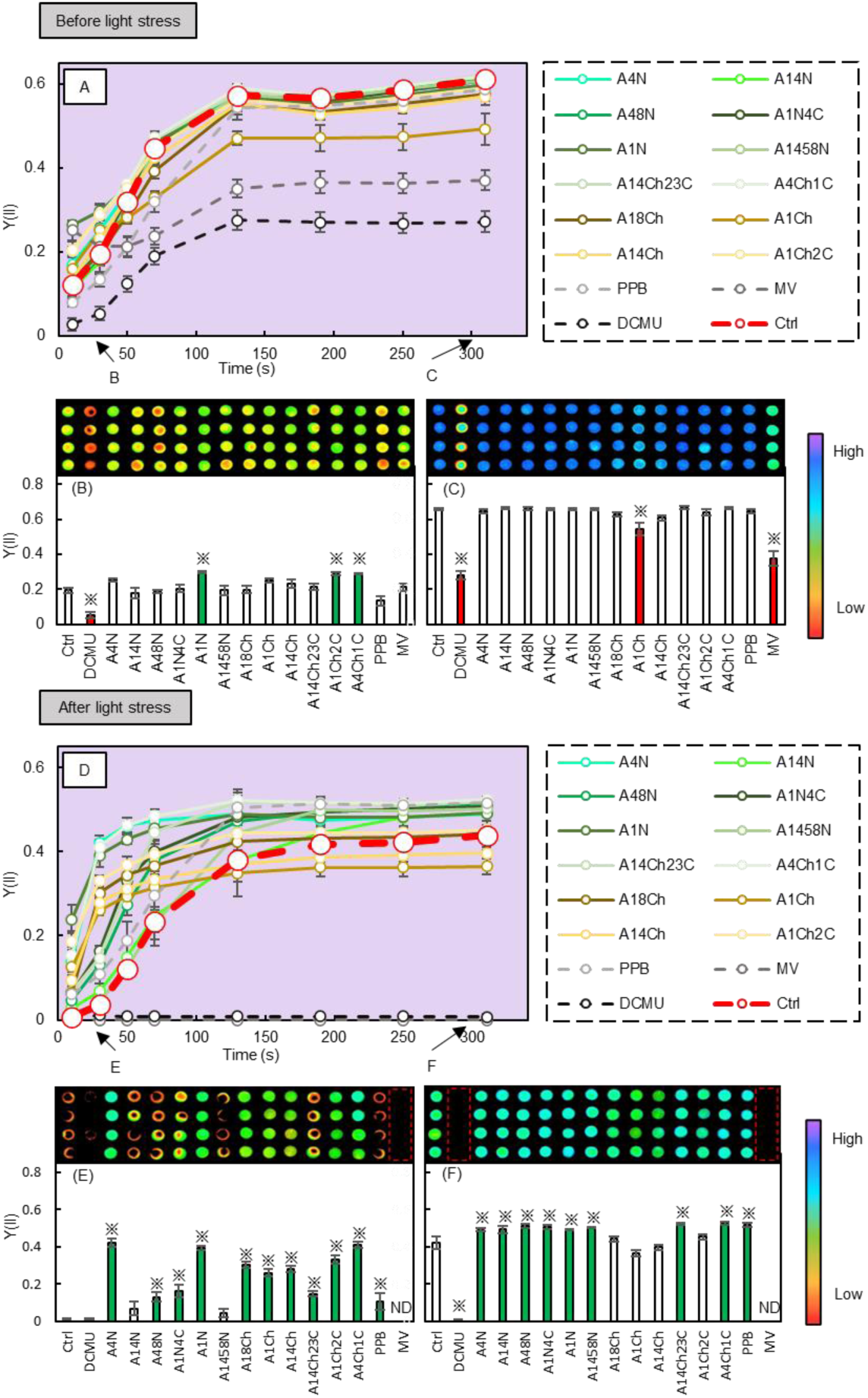
Photosynthesis induction curve of chemically treated leaf disks before and after light stress treatment. **A**. Photosynthesis induction curve of chemically treated leaf disks before light stress treatment, with actinic light be 200 µmol m^-2^ s^-1^. The arrows on the left and right side indicate data shown in **B** and **C**. The bold dotted line is the control group. **B, C**. Y(II) of the chemically treated leaf disks at the initial and steady state of photosynthesis induction before light stress treatment. **D**. Photosynthesis induction curve of chemically treated leaf disks after 12 h of light stress treatment, with actinic light be 200 µmol m^-2^ s^-1^. The arrows on the left and right side indicate data shown in **E** and **F**. The bold dotted line is the control group. **E, F**. Y(II) of chemically treated leaf disks at the initial and steady state of photosynthesis induction, after light stress treatment. The pseudo-color Y(II) image of **B, C, E, and F** are placed at the top of each corresponding chart. **Abbr.:** Ctrl: control, PPB: phenyl-p-benzoquinone, MV: methyl viologen, ND: not determined. Data is mean ± SE. ※ p<0.05, the red color indicates means that are significantly lower than the control group. The green color indicates means that are significantly higher than the control group *n* = 6.

The Y(II) of different groups before and after light stress treatment was also determined. The Y(II) of leaf disks adapted to dark environments often reaches its plateau value after an initial induction period from only several seconds to minutes after being exposed to a certain light intensity, which is called photosynthetic induction. In this study, photosynthetic induction of leaf disks was taken as a parameter to evaluate the effect of chemicals effect before and after light stress treatment. Y(II) at the initial state of photosynthetic induction (the first 30 seconds) and the steady state of photosynthesis (by the end of 310 seconds) were analyzed (Figure 2A). At the initial state of photosynthetic induction, A1N, A1Ch2C, and A4Ch1C treatment groups showed significantly higher Y(II) than the control group, while the Y(II) of the DCMU treatment group was significantly lower than the control group (Figure 2B). The A1N, A1Ch, and A14Ch treatment groups showed slightly higher Y(NPQ) compared with the control group, and the Y(NPQ) of the DCMU treatment group was lower than the control group (Figure S1C). The A1N, A1Ch, A1Ch2C, and A4Ch1C treatment groups showed slightly higher qP and the DCMU treatment groups showed lower qP compared with the control group Figure S1E). At the steady state of photosynthesis, the Y(II) of most chemical treatment groups showed no significant difference with the control group except for the A1Ch treatment group, which had significantly lower Y(II) than the control group (Figure 2C). The Y(II) of the DCMU and MV treatment groups were also significantly lower than that of the control group due to PSII inhibition and cell damage (Figure 2C). No significant difference in Y(NPQ) was observed except for the DCMU and MV treatment groups, which were significantly higher than the control group (Figure S1D). Also, no significant difference in qP was observed except for the DCMU treatment group, which was significantly lower than the control group (Figure S1F).

After light stress treatment for 12 hours, photosynthetic induction of the control group appeared to be slower than most of the chemical treatment groups, while photosynthesis in the DCMU and MV treatment groups was almost undetectable (Figure 2D). At the initial state of photosynthetic induction, A4N, A48N, A1N4C, A1N, A18Ch, A1Ch, A14Ch, A14Ch32C, A1Ch2C, and A4Ch1C treatment groups showed significantly higher Y(II) compared with the control group (Figure 2E), and at the steady state of photosynthesis, A4N, A14N, A48N, A1N4C, A1N, A1458N, A14Ch23C, and A4Ch1C treatment groups showed significantly higher Y(II) compared with the control group (Figure 2F). Notably, the PPB treatment group showed significantly higher Y(II) at both the initial and steady state of photosynthesis, indicating a minor protective effect against light stress (Figure 2E and 2F). In general, A4N, A48N, A1N4C, A1N, A14Ch23C and A4Ch1C treatment groups showed significantly higher Y(II) compared with the control group throughout photosynthetic induction. The A18Ch, A1Ch, A14Ch, and A1Ch2C treatment groups showed significantly higher Y(II) at the initial state but not at the steady state of photosynthesis. The A14N and A1458N treatment groups showed significantly higher Y(II) at the steady state of photosynthesis, but no significant difference in these two groups was observed at the initial state of photosynthetic induction. At the initial state of photosynthetic induction after light stress treatment, A4N, A1N, A18Ch, A1Ch, A14Ch, A1Ch2C, and A4Ch1C treatment groups showed significantly lower Y(NPQ) compared with the control group, while the Y(NPQ) of the MV treatment group was not detectable (Figure S1G). The A4N, A48N, A1N4C, A1N, A18Ch, A1Ch, A14Ch23C, A1Ch2C, and A4Ch1C treatment groups showed significantly higher qP compared with the control group, whereas the qP of the A4N, A1N, A18Ch, A1Ch, A1Ch2C, and A4Ch1C treatment groups was more than eight times higher than the control group (Figure S1I). During the steady state of photosynthesis after light stress treatment, A4N, A48N, A1N, A18Ch, A14Ch, A14Ch23C, A1Ch2C, and A4Ch1C treatment groups showed significantly lower Y(NPQ) compared with the control group, while the Y(NPQ) of the MV treatment group was not detectable (Figure S1H). The A1Ch and A14Ch treatment groups showed significantly lower qP compared with the control group, while the qP of the MV treatment group was not detectable (Figure S1J).

In order to further confirm the relationship between chemical treatment and the improvement of photosynthesis after light stress treatment, we studied the concentration-dependency of the protective effect of A1N and A4N because these two chemicals showed significantly higher Y(II) at both the initial and steady state of photosynthesis after light stress treatment (Figure 2E and 2F), and they are also similar in structure (Figure 1C). We also used Anthraquinone (9,10-anthraquinone, ANTQ) for treatments to see if similar effects could be observed with the skeletal structure alone. The chemicals were applied at different concentrations. And after exposure to light stress at 700 μmol photons m^−2^ s^−1^, the Y(II) of leaf disks treated with all three chemicals showed improved photosynthetic induction at all concentrations compared with the control group (Figure 3A, 3B, and 3C). It was also observed that A1N and A4N at high concentrations (4 μg ml^-1^) would cause significant decreases in Fv/Fm after light stress treatment (Figure 3D), but leaf disks treated with ANTQ showed significantly lower Fv/Fm at most concentrations except for 0.08 μg/ml (Figure 3D). The T_50_ (the time to reach half of the maximum value) of Y(II) in photosynthetic induction showed obvious concentration-dependence behavior in which the Y(II) of the leaf disks treated with high concentrations of chemicals reached plateaus faster than those treated with low concentrations (Figure 3E). It was also observed that leaf disks treated with A1N showed faster photosynthetic induction at lower concentrations compared with A4N and ANTQ (Figure 3F).

**Figure 3.**
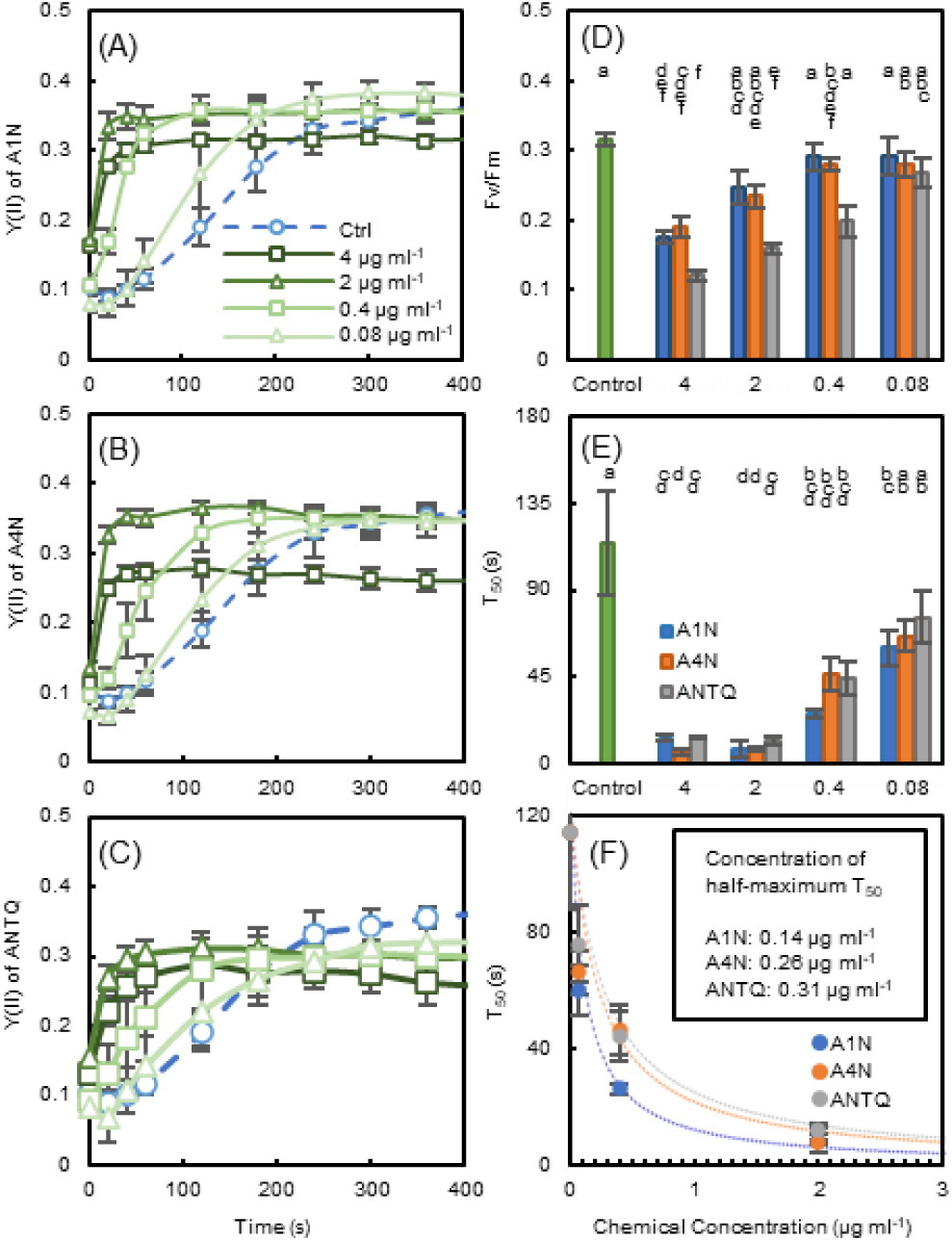
Photosynthesis inductions of leaf disks treated with various chemicals with different concentrations after 12 h light stress treatment. **A, B, C**. Photosynthesis inductions of leaf disks treated with A1N, A4N and anthraquinone (ANTQ) with different concentrations after exposure to high light conditions, with actinic light be 200 µmol m^-2^ s^-1^. **D**. Fv/Fm of leaf disks treated with various chemicals with different concentrations after exposure to high light conditions. **E, F**. T_50_ of Y(II) in photosynthetic inductions of leaf disks treated with various chemicals with different concentrations after exposure to high light conditions. **Abbr.**: Ctrl: control, ANTQ: anthraquinone. Data is mean ± SE. Bars with the same letter are not significantly different *n* = 4.

### Electron accepting abilities of A1N and A4N

Due to the quinone structure of the 9,10-anthraquinone backbone and the similar results of the chemicals groups and the PPB group for photosynthetic induction, it is possible that these chemicals function by serving as electron acceptors at certain steps of the photosynthetic electron transport chain. Therefore, we determined the electron accepting ability of A1N and A4N in the photosynthetic electron transport chain because they showed significantly higher Y(II) at both the initial and steady state of photosynthesis during induction after light stress treatment (Figure 2E and 2F) and because they also had similar structures (Figure 1C). In order to evaluate the electron accepting ability of A1N and A4N, we replaced the PPB or MV with chemicals with the same concentration so that the electron transport rate with chemicals or PPB/MV serving as electron acceptors could be compared.

For measurements of the whole chain electron transport rate, where electron transport from H_2_O to the donor side of PSI was measured, A1N showed a similar electron accepting ability as MV, while A4N showed lower electron accepting ability than MV and A1N (68% of MV) (Figure 4A). This result also indicated that, based on the principles of this measurement described by Yamasaki et al. (2002), A1N and A4N may react with O_2_ upon being reduced, which is similar to MV. However, electron accepting by A1N and A4N could happen at any point between PSII and PSI. In order to identify the exact location of electron accepting, we further measured electron transport at PSII and PSI separately. In measurements of electron transport at PSII, where electron was transported from H_2_O to the acceptor side of PSII, electron accepting ability of A1N and A4N was not detected (Figure 4B). In measurements of electron transport at PSI, where the electron was transported from DCPIP to the acceptor side of PSI, A1N and A4N showed lower but obvious electron accepting abilities compared with MV (A1N was 86% and A4N was 78% of MV) (Figure 4C). These results indicate that A1N and A4N could serve at least partially as electron acceptors at PSI, but not at PSII.

**Figure 4.**
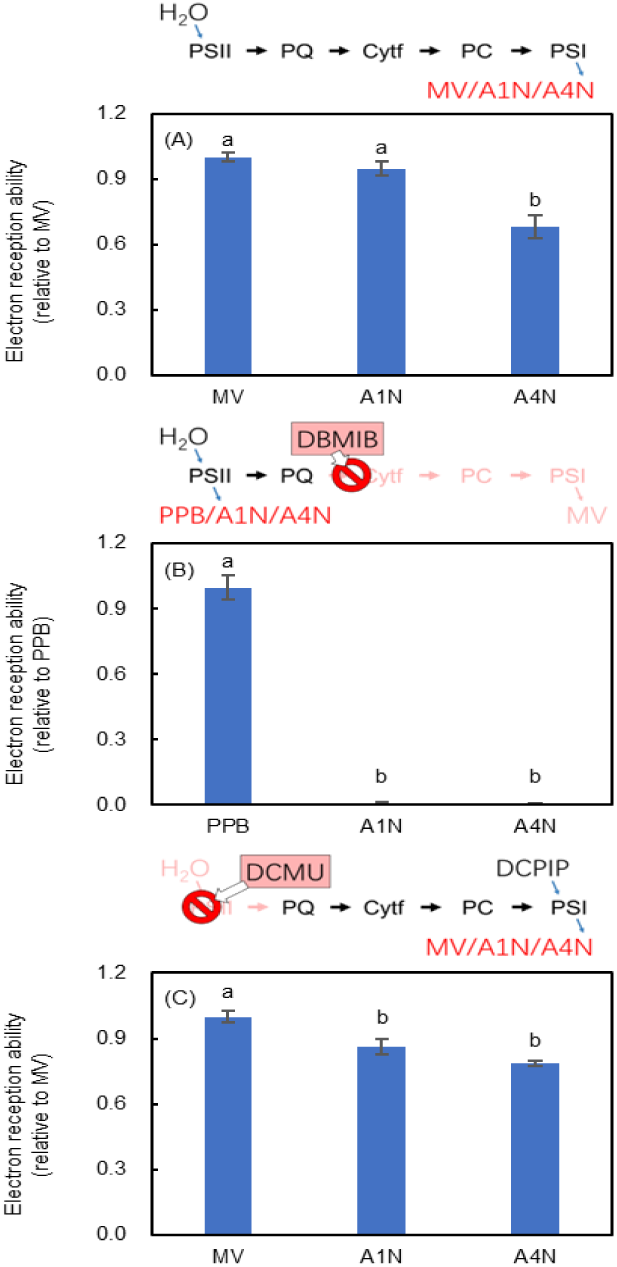
Electron reception ability of A1N and A4N compared with methyl viologen (MV) and phenyl-p-benzoquinone (PPB) in whole chain PSII and PSI electron transport measurements. **A**. The electron reception ability of A1N and A4N is compared with MV in the whole chain electron transport measurement. The electron reception of MV is normalized as 1. **B**. The electron reception ability of A1N and A4N is compared with PPB during PSII electron transport measurement. The electron reception of PPB is normalized as 1. **C**. The electron reception ability of A1N and A4N is compared with MV during PSI electron transport measurement. The electron reception of MV is normalized as 1. The electron transport pathway of each measurement is indicated above in the corresponding chart. Data is mean ± SE. Means with different letters are significantly different *n* = 4.

### The short-term effect of chemicals on Photosynthetic characteristics in intact leaf of tobacco

In this study, the effect of chemicals on intact tobacco leaves was investigated. A1N was chosen for chemical treatment due to its high electron accepting ability and the high Y(II) value in photosynthetic induction. After light stress treatment, both the control group and the A1N treatment group showed a decrease in Fv/Fm, but no significant difference between the control group and the A1N treatment group was observed at each measurement time point (Figure 5A). Chlorophyll fluorescence was also determined simultaneously with gas exchange. The ETR II and CO_2_ assimilation rates showed declines after light stress treatment in the control group and the A1N treatment group, but the decrease was significantly more pronounced in the control group (Figure 5B and 5C). The control group showed significantly lower ETR II, especially at 72 hours of light stress treatment, while the ETR II of the A1N treatment group showed no significant changes throughout the light stress treatment (Figure 5B). The control group showed a significantly lower CO_2_ assimilation rate at 72 hours of light stress treatment, while the CO_2_ assimilation rate of the A1N treatment group showed no significant change throughout the light stress treatment (Figure 5C). In order to investigate whether the variations in ETR II and CO_2_ assimilation rates were caused by variations in CO_2_ transport rate involved in stomatal gas exchange, we further analyzed the stomatal conductance and intercellular CO_2_ concentrations. Declines in stomatal conductance and intercellular CO_2_ concentrations after light stress treatment were observed in the control group and the A1N treatment group, but no significant difference was observed between them at each measurement time point (Figure S2A, S2B, S2C, and S2D). In contrast, tobacco leaves treated with A1N showed no significant difference in Fv/Fm and CO_2_ assimilation rates compared with the control group with the absence of strong light treatment (Figure S3). After 72 hours of light stress treatment, decreases in leaf greenness, especially in the control group, were observed (Figure 5E), and the chlorophyll concentrations of the control group were significantly lower than the A1N treatment group (Figure 5D).

**Figure 5.**
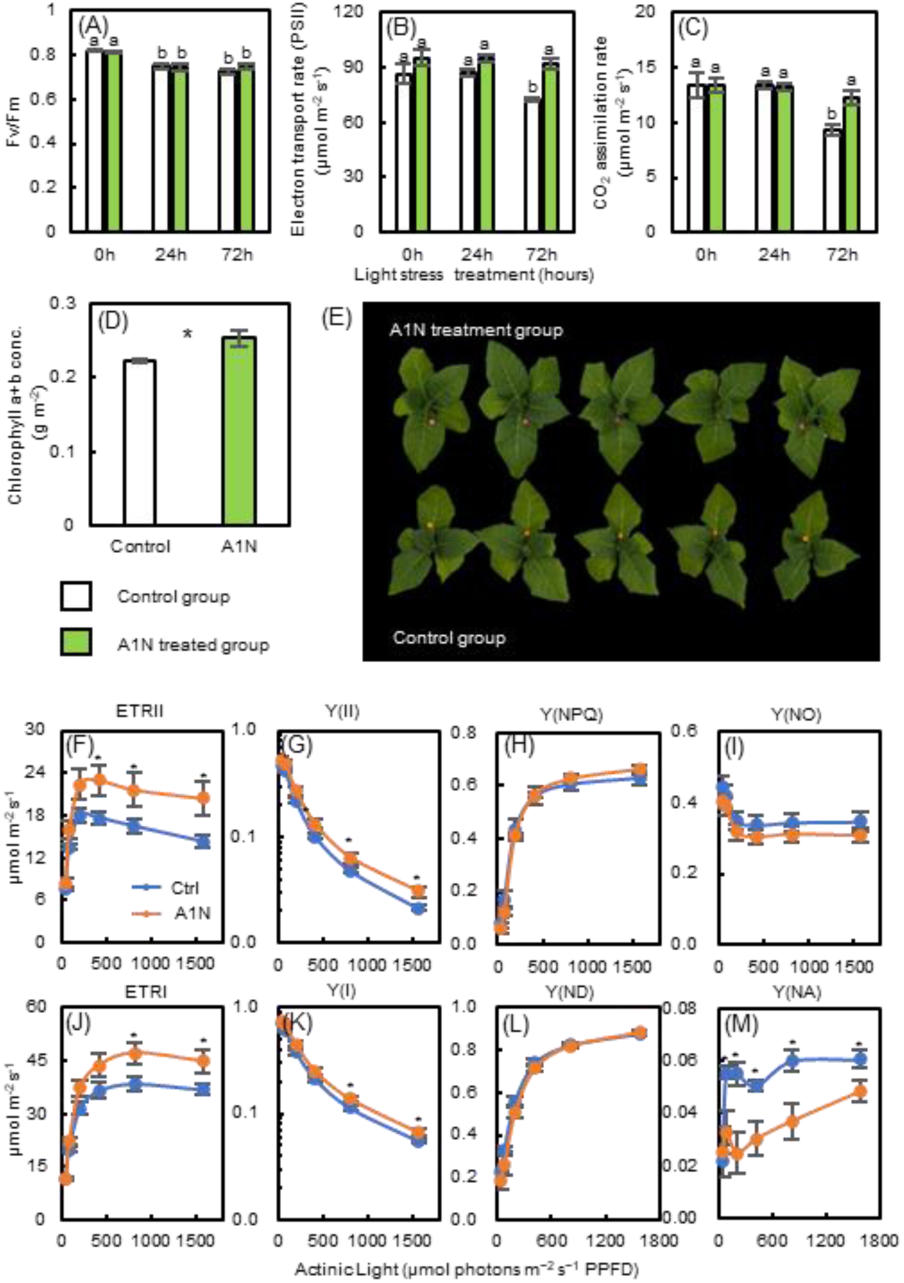
Photosynthesis parameters of tobacco leaves after light stress treatment. **A, B, C.** Fv/Fm, Electron transport rate of PSII, and CO2 assimilation rate of tobacco leaves after 0 h, 24 h, and 72 h of light stress treatment. **D.** Concentration of chlorophyll a+b after 72 h of light stress treatment. **E.** Tobacco plants after 72 h of light stress treatment are shown. The leaves of the A1N group were marked with pink stickers, and the leaves of the control group were marked with yellow stickers. **F, G, H, I.** Electron transport rate, Y(II), Y(NPQ), Y(NO) of PSII in tobacco leaves after 72 h of light stress treatment. **J, K, L, M.** Electron transport rate, Y(I), Y(ND), Y(NA) of PSI in tobacco leaves after 72 h of light stress treatment. **Abbr.**: Ctrl: control. Data is mean ± SE. Means with different letters are significantly different. ns p>0.05, * p<0.05, *n* = 5.

A Dual-PAM-100 was used to investigate the impact of light stress on PSII and PSI in tobacco leaves. After exposure to light stress at 700 μmol photons m^−2^ s^−1^ for 72 hours, the ETR II and Y(II) of the A1N treatment group were significantly higher than the control group (Figure 5F and 5G), especially for high light conditions (400– 1500 μmol photons m^−2^ s^−1^). However, the Y(NPQ) and Y(NO) of the control group and the A1N treatment group showed only minor differences (Figure 5H and 5I). For PSI activities, the ETR I and Y(I) of the A1N treatment group were significantly higher than the control group at high light conditions (higher than 800 μmol photons m^−2^ s^−1^) (Figure 5J and 5K). Such differences could be explained by significantly lower Y(NA) in the A1N treatment group (Figure 5M). However there was no difference between the A1N treatment group and the control group for Y(ND) (Figure 5L).

### The long-term effects of chemicals on the photosynthetic and growth characteristics of Arabidopsis and lettuce

In order to investigate the long-term effects of chemicals on plant recovery and growth after light stress treatment, the effects of the chemicals on whole Arabidopsis and lettuce plants were investigated. For Arabidopsis, although no significant differences in Fv/Fm was observed during this stage, the A1N treatment group showed higher Y(II) and lower Y(NPQ) and Y(NO) compared with the control group after 96 hours of light stress treatment and actinic light at cultivation levels and strong light stress levels (Figure 6A and 6B). Similar results were observed for lettuce as well. The A1N treatment group showed higher Fv/Fm after 96 hours of light stress treatment (Figure 7A and 7B) and actinic light at cultivation levels. It also showed higher Y(II) (Figure 7A and 7B) with actinic light at strong light stress levels and higher Y(II) and lower Y(NO) (Figure 7A and 7B) compared with the control group. Moreover, the A1N treatment group showed higher accumulation of biomass by the end of the light stress treatment (Figure 7 C).

**Figure 6.**
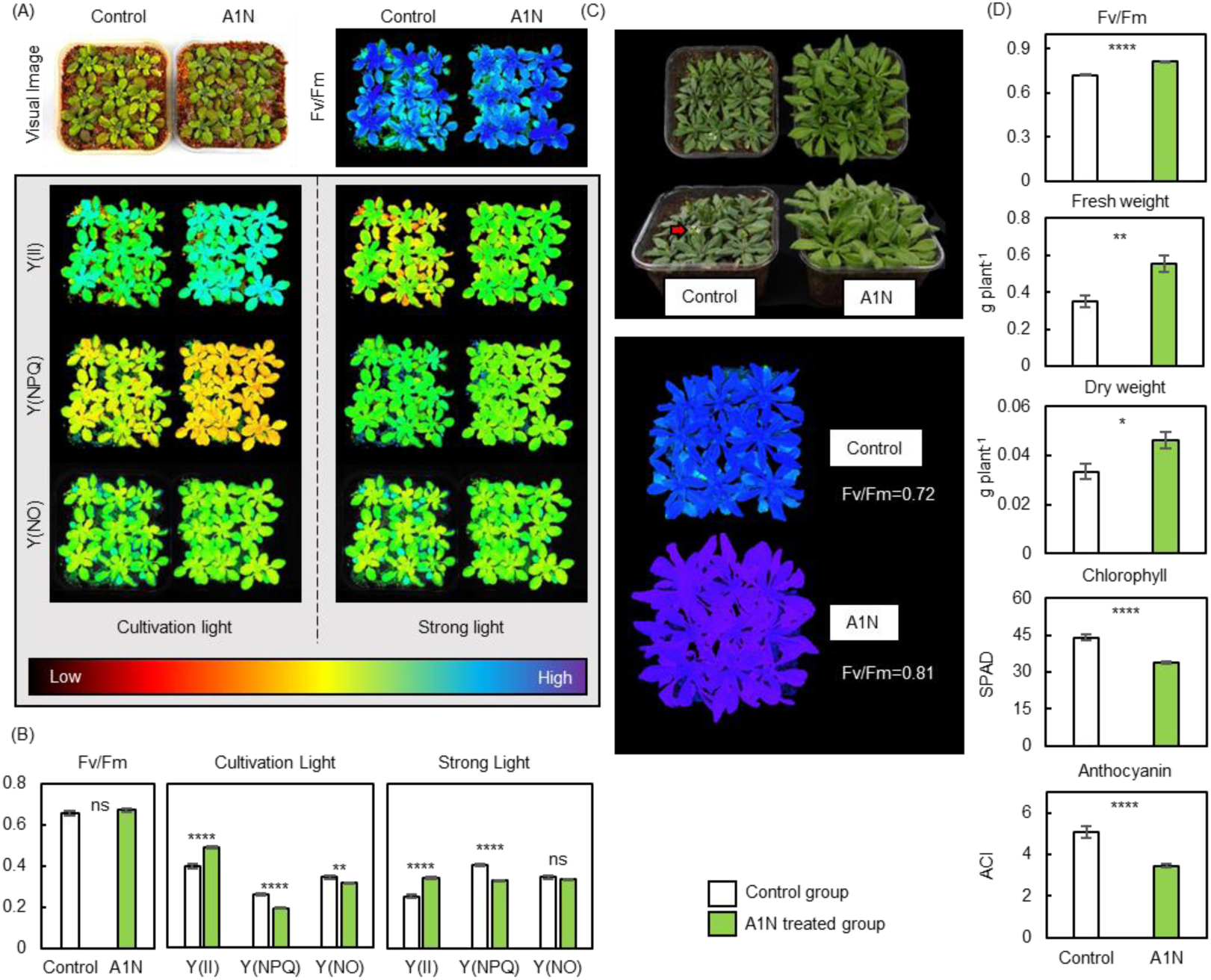
Arabidopsis after exposure to light stress for 96 hours and 1 week’s recovery. **A**, **B**. The visual images, pseudo-color (indexed color mode) images and data of Fv/Fm, Y(II), Y(NPQ), and Y(NO) of plant exposed to light stress for 96 hours, under actinic light at cultivation levels (200 μmol photons m^−2^ s^−1^) and strong light stress levels (700 μmol photons m^−2^ s^−1^). **C**. The visual images and pseudo-color (indexed color mode) images of the Fv/Fm of plants after 1 week’s recovery. The red arrow points to early bloating. **D**. Fv/Fm, above ground fresh weight, above ground dry weight, the chlorophyll and anthocyanin contents of plants after 1 week’s recovery. Data is mean ± SE. ns p>0.05, * p<0.05, ** p<0.01, **** p<0.0001, *n* = 9.

**Figure 7.**
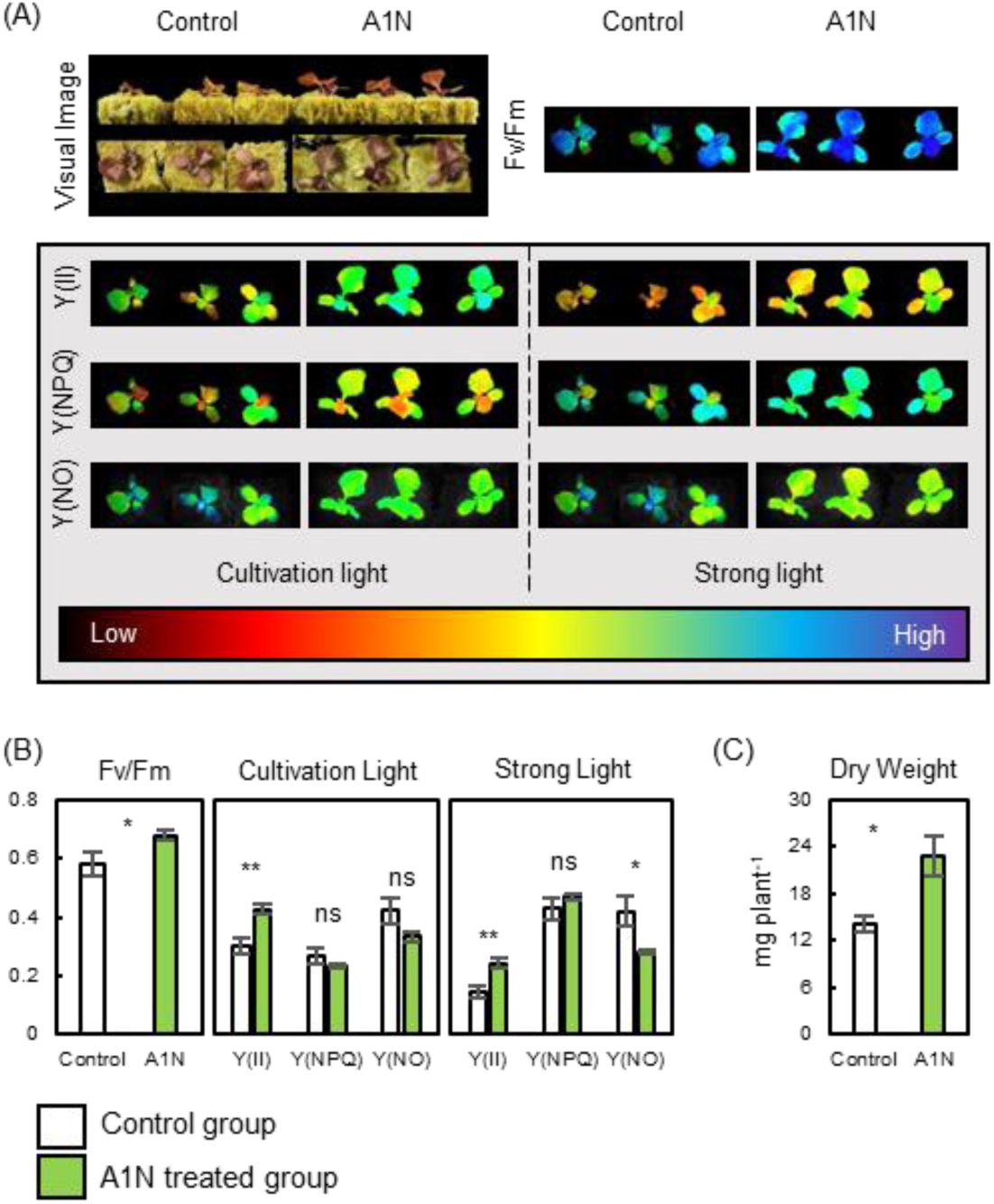
Lettuce after exposure to light stress for 96 hours. **A**. **B**. Visual images of plants, pseudo-color (indexed color mode) images and data of Fv/Fm, Y(II), Y(NPQ), and Y(NO) under actinic light at cultivation levels (200 μmol photons m^−2^ s^−1^) and strong light stress levels (700 μmol photons m^−2^ s^−1^) are displayed, respectively. **C**. Above ground dry weight after exposure to light stress for 96 hours. ns p>0.05, * p<0.05, ** p<0.01, *n* = 6.

To investigate the effect of the A1N treatment on global transcriptional responses toward high light stress in whole Arabidopsis plants, RNA-seq analysis was conducted. A total of 76 genes were identified as differentially expressed genes (DEGs) between the A1N treatment group and the control group, of which 37 genes were up-regulated and 39 genes were down-regulated in the control group (Table S1). GO enrichment analysis of the DEGs showed that several GO terms that were related to photosynthesis were enriched (Figure S6A). Therefore, we further examined the expression of the gene encoding photosynthesis apparatus and carbon fixation (Figure S6B, Table S2). Some genes related to the light-harvesting chlorophyll protein complex, the electron transport chain, and the carbon fixation were slightly down-regulated in the control group compared with the A1N treatment group.

After the light stress treatment, the Arabidopsis plants were restored to the condition of non-stressful cultivation. After 1 week’s recovery, plants in the control group showed inhibited growth and early flowering, but plants in the A1N treatment group and the group that was not treated with light stress showed almost no sign of stress (Figure 6C). The Fv/Fm of the control group was 13% lower than the A1N treatment group and 11% lower than the group that was not treated (Figure 6C and 6D). The fresh weight and the dry weight of the control group were also significantly lower than those in the A1N treatment group and in the group that was not treated (Figure 6D). Furthermore, plants in the control group showed significantly higher chlorophyll and anthocyanin concentrations and deeper colors than the A1N treatment group (Figure 6C).

## Discussion

By performing chemical screening with a novel system (Figure 1A), several anthraquinone derivatives with protective effects against light stress were discovered (Figure 1C). Tobacco leaf disks, intact leaves, and whole Arabidopsis plants treated with the chemical A1N showed improved photosynthesis and plant growth during and after light stress treatment (Figure 2, 3, 5, and 6). The electron accepting ability around the PSI was observed for A1N and A4N (Figure 4). Therefore, the protective effect of these anthraquinone derivatives can be explained by their ability to remove excess electrons from PSI. Moreover, it should be noted that the A1N treatment showed no negative effect on plants in non-stress conditions, and intact tobacco leaves treated with A1N showed no significant difference in photosynthesis compared with the control group when continually kept in cultivation conditions (Figure S3).

### Anthraquinone derivatives may serve as electron acceptors

Anthraquinone derivatives are groups of chemicals with robust electrochemical and photochemical activities (Diaz, 1990; Uchimiya and Stone, 2009). It was noticed that phenyl-p-benzoquinone, which is an electron acceptor on the donor side of PSII, showed similar but lower protective effects for leaf disks after light stress (Figure 2). Therefore, it is reasonable to suspect that the protective effect of anthraquinone derivatives is related to their electron accepting ability as quinones. In order to prove this, electron transport in the isolated thylakoid membrane was measured with an oxygen electrode-based determination system. According to the results of the oxygen electrode, the chemicals A1N and A4N showed similar electron accepting abilities compared to MV around PSI but not PSII (Figure 4). This is also explained by the differences in the reduction potentials of A1N, PSI, and PSII. According to previous studies, the reduction potential of A1N is -816 mV vs. The standard hydrogen electrode (*E*_1/2_ refers to the first reduction feature) (Gallmetzer et al., 2022) is lower than MV (-446 mV) but higher than the reduced P700 (-1320 mV) during PSI (Kim et al., 2009; Khorobrykh et al., 2020) (Figure S4). However, the redox potential of PSII is -620 ∼ -660 mV (Kato et al., 2009; Khorobrykh et al., 2020), which is too low for A1N reduction (Figure S4). These results indicate that A1N can be reduced around PSI primary electron acceptors (A_0_ and A_1_) (Figure S4) upon light illumination but not during PSII. Therefore, it is possible that A1N and other anthraquinone derivatives are engaged in the removal of excess electrons from PSI under light stress conditions. This is further corroborated by the increased qP at the initial state of photosynthetic induction before and after light stress treatment (Figure S1E, I) and decreased Y(NA) during PSI after light stress treatment (Figure 5M).

It should be noted that the reduction potential of anthraquinone derivatives is largely influenced by other residues connected to the backbone. For A1N, a NH_2_-residue is connected to the anthraquinone backbone (Figure 1C). NH_2_-is an electron-donating residue that decreases the reduction potential of A1N by approximately 132 mV compared with anthraquinone (-684 mV) (Gallmetzer et al., 2022). A1N and A4N have a single electron-donating residue (Figure 1C) and have showed strong protective effects against light stress in leaf disks (Figure 2 and 3). However, derivatives with more than one electron-donating residue, such as A14N, A48N, and A14N58N, showed relatively lower protective effects compared with A1N and A4N (Figure 2), indicating that overly decreasing the reduction potential reduce can have a negative effect.

Although we confirmed that A1N and A4N could serve as electron acceptors in photosynthetic electron transport, details about the electron accepting process are not clear as the electron accepting ability of A1N and A4N in whole chain measurements did not perfectly match with the PSI measurements (Figure 4A and 4C). However, it is possible that the final electron sinks for A1N and A4N mediated electron transport is O_2_ as they showed strong electron acceptance in whole chain and PSI electron transport measurements. This was also the case with MV, which reacted with O_2_ upon electron acceptance and reduced O_2_ into superoxide. Therefore, electron transport can be detected by investigating the O_2_ consumption rate in the solution. As a side effect, MV can cause drastic cell death due to the accumulation of toxic superoxide especially around the iron-sulfur center of PSI, inducting Fenton reaction which could damage the iron-sulfur center of PSI (Sonoike 1995). However, it is not surprising that this effect turned out to be absent in most anthraquinone derivatives we studied. This is because the reaction between anthraquinone derivatives and O_2_ upon reduction is slightly different with MV. The reaction of anthraquinone, electrons, and O_2_ lead to the formation of endoperoxo complex intermediates rather than free radicals like superoxide, and these intermediate compounds eventually turn into H_2_O_2_ (Nishimi et al. 2011) possibly at more distant locations from the iron-sulfur center of PSI (Figure S4). Therefore, although O_2_ serves as a main electron sink for anthraquinone derivatives, this process can be less lethal than reactions between other electron acceptors, such as MV or O_2_ only, because H_2_O_2_ around PSI iron-sulfur center could be much lower.

### Anthraquinone derivatives improve photosynthesis and growth during and after light stress

In leaf disks, improved photosynthetic induction and higher initial state qP were observed before and after light stress treatment (Figure 2 and S1), indicating that these chemicals could improve the quantum yield of PSII by providing a higher oxidation state of the primary donor P700 of PSI at the very beginning of photosynthetic induction. For intact tobacco leaves, photosynthetic activities were well preserved throughout light stress treatment in the A1N treatment group, but they decreased significantly in the control group (Figure 5). Moreover, such changes in photosynthesis are not likely to be induced by CO_2_ concentrations. Although light induced stomatal closure was observed in light stress treatments, no significant differences in intercellular CO_2_ concentrations and stomatal conductance were observed between the control group and the A1N treatment group (Figure S2). For whole Arabidopsis and lettuce plants, A1N-treated plants showed higher Fv/Fm and Y(II) and lower Y(NPQ) and Y(NO) with better recovery and fewer signs of stress (Figure 6 and 7), while the control group showed lower Fv/Fm and higher accumulation of chlorophyll and anthocyanin together with inhibited growth and early flowering, which is considered to be a sign of stress factors (Figure 6) (Wada and Takeno, 2010). It has been reported that the genes encoding photosynthesis apparatus are repressed during high light stress to mitigate excess absorbed light (Huang et al., 2019). In this study, A1N-treated plants showed suppression of some photosynthesis-related gene expressions compared with the control group (Supplementary Fig. A). These results suggest A1N maintained photosynthesis capacity during high light stress by protecting the photosynthetic apparatus from excess light in the transcription levels.

Previous studies have shown that some anthraquinone derivatives may serve as a PSII inhibitor by competing over the Q_B_ site on D1-proteins due to their similarity with plastoquinone and that this causes negative effects in electron transport and plant photosynthesis in the short term (Strotmann et al., 1982; Oettmeier *et al*., 1988). However, no significant decrease in Fv/Fm was observed in most of the chemical treatment groups for the leaf disks (Figure S1A and S1B) and tobacco leaves (Figure 5A and S3A) with or without light stress treatment, and it was only observed in A1N and A4N when the concentrations were very high (4 μg ml^-1^) (Figure 3D). This difference is possibly caused by variations in treatment concentrations. In studies mentioned above, anthraquinone derivatives up to 50 μM were applied directly onto separated thylakoid membranes, while in our study, 20∼40 μM (2∼4 μg ml^-1^) of anthraquinone derivatives were applied to the outer surface of plant materials and left to diffuse into chloroplasts naturally. Therefore, although the treatment concentrations in studies by Strotmann et al. (1982) (I_50_ = 25 μM) were similar with those in this study for tobacco leaf disks, and intact leaves, the final concentrations of anthraquinone derivatives presented in chloroplast could be two orders lower in the case of this study (∼0.2 μM). When high concentrations (50 μM) of A4N was applied together with electron acceptors such as MV or PPB directly to thylakoid membranes to evaluate their PSII inhibition effects, significant decreases in thylakoid membrane electron transport rates at whole chain and PSII, but not in PSI, were observed (Figure S5A and S5B). These results showed that beside with serving as electron acceptors, high concentrations of anthraquinone derivatives can also significantly inhibit PSII, as indicated by Shortamnn et al. (1982). With lower concentrations, A1N and several other anthraquinone derivatives can protect plants from light stress, improve photosynthesis during light stress, and result in better post-stress recovery, but at higher concentrations, they can also inhibit PSII functions.

Treating plants with certain chemicals to achieve higher tolerances against stresses is a well-explored strategy with many exciting achievements. Over the past decades, there has been increasing evidence that plants can be sensitized for more rapid or more intense mobilization of resistance systems, leading to enhanced tolerance against biotic and abiotic stresses (Conrath et al. 2006). The physiological state in which plants are able to activate defense responses faster or better is called the primed state of the plant. The use of chemicals to induce the primed state is an interesting topic not only as a mechanism for fundamental research but also for practical applications (Jakab et al., 2005; Kim et al., 2017). However, the engagement of stress-response genes in primed state plants results in inhibited growth or reduced photosynthesis, so there is no benefit if there is no stressor (Martinez-Medina et al., 2016). Therefore, a precise prediction of stress conditions is necessary, and plants have to be treated at the right time before the stressor to fully benefit from the priming effect, which is very difficult to achieve in practical applications. But in this study, no decrease in photosynthesis was observed in A1N-treated tobacco in non-stressful light conditions (Figure S3). This indicates that we can treat plants with chemicals, such as A1N, to prevent light stress at any time without worrying about the negative effects.

### Anthraquinone derivatives can be used as agrochemicals

Anthraquinone derivatives are widely found in natural environments. In lichen, they have been found to provide protection against UV-B damage (Nybakken *et al*., 2004), and they are present in rhizomes, flowers, and fruits. More than 200 compounds belonging to these derivatives were identified in plants. They play various roles, such as the regulation of energy transfer and cell death (Diaz-Munoz et al., 2018). Due to their wide range of colors, anthraquinone derivatives are used as dyes in industry (Routoula and Patwardhan, 2020). They are also well known for their anticancer, anti-inflammatory, diuretic, anti-arthritic, antifungal, antibacterial, and antimalarial properties (Diaz-Munoz et al., 2018). They are also used in many traditional herbal medicines to cure digestive system disorders (Wang *et al*., 2008). Some anthraquinone derivatives have been used for nonlethal pest management in agricultural production since the 1940s (DeLiberto and Werner, 2016). In organisms, there are two major pathways that are engaged in the synthesis of anthraquinone chemicals. The first is the polyketide pathway, which is observed in *Photorhabdus luminescens*, a gammaproteobacterium of the family Morganellaceae. It is a lethal pathogen of insects and higher plants, including *Senna tora* (Brachmann et al., 2007; Kang et al., 2022). In this pathway, octaketide is dehydrated, and several anthraquinone derivatives, including Endocrocin, Physcion, Rhein, and Chryso-obtusin, are synthesized (Kang et al., 2022). The second pathway is a combination of shikimate and mevalonate/methyl-D-erythritol 4-phosphate pathways, which is observed in *Cinchona officinalis* and *Rheum tanguticum* (Kang et al., 2020; Zhou et al., 2021). In this pathway, products from the Shikimate and Terpenoid pathways, including shikimic acid, glucose, acetyl CoA, pyruvate, and glyceraldehye 3-P, are further processed to synthesize anthraquinone derivatives (Kang et al., 2020). Although these two pathways have not been found in Arabidopsis or tobacco, it would be an interesting avenue of research to verify whether these two pathways for anthraquinone biosynthesis are also functional in Arabidopsis, tobacco, and other crop plants and whether it is possible to introduce anthraquinone synthesis pathways into these plants. If it is, it could result in the natural accumulation of such chemicals within plant cells and provide them with endogenous protection against light stress. However, the detailed biosynthesis of anthraquinone is still unclear for plant species. According to previous studies in *Rheum tanguticum* and *Senna tora*, a total of more than 200 candidate enzyme genes are involved in the synthesis of anthraquinone (Kang et al., 2020; Zhou et al., 2021), and it is hard to tell which of them are essential for the introduction of anthraquinone synthesis pathways in other plants. So further studies are required.

In this study, A1N showed significant protective effects against light stress in tobacco, Arabidopsis, and lettuce (Figure 2, 3, 5, 6, and 7) with no negative effects in non-stressful conditions (Figure S3). Moreover, there was no data suggesting that A1N is toxic. Therefore, A1N or A1N analogs are candidate compounds for the development of light stress protectors in agriculture. Utilization of these chemicals for enhanced light stress tolerance offers a new method for securing the productivity of food crops and the commercial benefits of high value products, including fruits, vegetables, flowers, herbs, and ornamental plants, under extreme light intensities. Hopefully, these findings will inspire the development of new techniques that enhance the tolerance of plants in order to address the challenges of this unprecedented period of global climate change.

## Materials and methods

### Chemical library and compounds

A total of 12,000 compounds from the chemical library provided by Maybridge were used in the screening together with 10 structural analogs, the photosynthesis inhibitor DCMU, the electron receptor phenyl-p-benzoquinone (PPB), and Methyl viologen (MV) (Figure 1C). Structural analogs were obtained from the database of ChemCupid. All chemicals were diluted with DMSO to a stock concentration of 2 mg ml^-1^ and kept at -80 °C.

### Plant materials and growth conditions

Tobacco (*Nicotiana tabacum* L. “Wisconsin-38”), Arabidopsis (*Arabidopsis thaliana* (L.) Heynh, “Columbia-0 (CS60000)”), and lettuce (*Lactuca sativa* [L.] “New Red Fire Lettuce”) were used in this study. The seeds of tobacco and Arabidopsis were sown in a 1:1 mixture of MetroMix (Metromix 350, Hyponex) and vermiculite. Lettuce was sown on rockwool. All plants were cultivated in a temperature-controlled growth chamber for the first 3–4 weeks. The growth chamber was operated with a daylight/dark period of 10/14 h, and PPFD in the daylight period was set to 150 μmol photons m^−2^ s^−1^. Air temperature and relative humidity were set to a constant 22 °C and 60%, respectively. Arabidopsis and lettuce cultivated in the growth chamber were used for experiments in the third week. After four weeks, tobacco plants were transplanted into φ252×300 mm pots with 100% Akadama soil (red granular soil) in a greenhouse, and 5 g of full nutrition fertilizer (A906, JCAM AGRI. CO., LTD.) was applied to each pot. Fully expanded tobacco leaves after 8–12 weeks (before flowering stage) of cultivation were harvested for further measurements. The spinach (*Spinacia oleracea*) used in this study was purchased at a local retail store.

### Construction of a chemical screening system using tobacco leaf disks

To ensure highly efficient screening of large amounts of chemicals, a system of 96 well plates loaded with leaf disks in each well was constructed (Figure 1A). Tobacco plants cultivated for 8–12 weeks were moved to a dark room 24 hours before the experiment. Leaf disks were collected from fully expanded leaves with a leaf punch (φ = 5.35 mm). To ensure the homogeneity of photosynthesis properties, only healthy leaf tissue that was not adjacent to leaf veins or the edges of leaves was used. Collected leaf disks were kept in a 96 well plate container for treatment and measurement, and one 7 mm surgical cotton ball was placed in each well of the 96 well plates (Thermo Fisher scientific). Black plates were used to minimize light reflection and penetration, which could cause noise when taking measurements. 200 μl of distilled water was added to each well to create a moist environment. Leaf disks were placed on the cotton balls with the top side of the leaves facing upward. They were kept wet throughout this process (Figure 1A).

Chemical components were added to the well of each plate for a final concentration of 2 μg ml^-1^. Plates loaded with diluted chemicals and leaf disks were infiltrated for 5 min, and chlorophyll florescence was then determined with the Imaging-PAM M-Series (IMAGING-PAM; Walz, Effeltrich, Germany) to evaluate the instant effect of the chemicals on photosynthesis. The leaf disks in the plates were then exposed to high light intensity for 12 h; the plates were sealed with cling film to prevent evaporation and placed in a growth chamber operated with a constant light period; and PPFD was set to 700 μmol photons m^−2^ s^−1^. Air temperature and relative humidity was set to a constant 22 °C and 60%, respectively. After high light treatment, chlorophyll florescence was again determined with the Imaging-PAM to evaluate the protective effect of the chemicals on photosynthesis against light stress (Figure 1B).

Each chemical could be applied onto one leaf disk (high throughput but low accuracy) or multiple leaf disks (low throughput but high accuracy). In this study, high throughput screening was used for the first screening of the whole chemical library, and low throughput screening was used for the second screening to confirm the results obtained in the first screening. A fully expanded tobacco leaf (major-axis length of over 30 cm) harvested between 8–12 weeks can provide enough leaf disks for approximately four 96 well plates. In this study, eight plates were processed in one day. Therefore, around 700 different chemicals could be tested in a day.

### The short- and long-term effects of the chemicals on living plants

The short-term (from hours to days) and long-term effects (from days to week) of the chemicals on the plants were determined separately. For short-term chemical effects, intact leaves from tobacco seedlings cultivated for 3–4 weeks in the growth chamber were used. Tobacco leaves (not detached from the plant) were immersed in 4 μg ml^-1^ of chemicals plus 0.2% solvent DMSO (chemical treatment group) or 0.2% DMSO only (control group) for 1 hour and then returned to cultivation conditions and kept in the dark for 12 hours. For light stress treatments, plants were moved to a growth chamber with the same temperature and relative humidity as the cultivation conditions, but PPFD was at a constant 700 μmol photons m^−2^ s^−1^. Gas exchange, chlorophyll fluorescence, and P700 redox state in treated leaves were determined with the LI-6400XT (Li-COR, Lincoln, NE), the Imaging-PAM M-Series (Walz, Effeltrich, Germany), and the DUAL-PAM-100 (Walz, Effeltrich, Germany) throughout the light stress treatments. Treated leaves were harvested, and the chlorophyll concentrations of treated leaves were determined after 72 hours of light stress treatment.

To analyze the long-term effects of the chemicals, the seedlings of Arabidopsis and lettuce cultivated for 3 weeks were used. Plants were sprayed with 20 μg ml^-1^ of chemicals plus 1% solvent DMSO (chemical treatment group) or 1% DMSO only (control group). For light stress treatment, plants were moved to a growth chamber with the same temperature and relative humidity as the cultivation conditions, but PPFD was at a constant 700 μmol photons m^−2^ s^−1^. For Arabidopsis, plants were treated with stressful light conditions for 96 hours and then returned to cultivation conditions and allowed to recover for 1 week before harvesting. RNAseq was determined by the end of the light stress treatment, and chlorophyll fluorescence was determined with the Imaging-PAM throughout the light stress treatment and after 1 week’s recovery. The SPAD value, anthocyanin concentration, and above ground biomass were determined after 1 week’s recovery. For the lettuces, plants were treated with stressful light conditions for 96 hours and harvested. Chlorophyll fluorescence was determined by the Imaging-PAM throughout the light stress treatment, and the above ground biomass was determined by the end of the light stress treatment.

### Analysis of gas exchange, chlorophyll fluorescence, and P700 measurements

The determination of the electron transport rate during PSII and PSI was conducted with the Maxi-version of the Imaging-PAM M-Series (Walz, Effeltrich, Germany) and the DUAL-PAM-100 (Walz, Effeltrich, Germany). All plant materials used for determinations mentioned above were adapted to darkness for at least 30 minutes before measurement started. For the chemical screening system for leaf disks, actinic light was set to 64 µmol m^-2^ s^-1^ in the Imaging-PAM system. To ascertain the long-term effects of the chemicals on Arabidopsis and lettuce seedlings, actinic light was set to 200 µmol m^-2^ s^-1^ and 700 µmol m^-2^ s^-1^. The chlorophyll fluorescence from plant materials was selected by defining the areas of interests (AOI) with the ImagingWin Software (Walz), and pseudo-color (indexed color mode) images of tested material could be obtained. For the Dual-PAM-100 measurements, intact leaves were clamped in the 1.4 cm^2^ chamber of a gas exchange system (GFS-3000, Walz Germany). To measure the short-term effects of the chemicals on tobacco seedlings, actinic light was set to gradients ranging from 30 to 1600 µmol m^-2^ s^-1^. To obtain stable fluorescence from the samples, each state lasted for 180 seconds. The maximum potential photochemical quantum yield of photosystem II (PSII) (Fv/Fm), the effective quantum yield of photochemical energy conversion during PSII (Y[II]), the quantum yield of regulated energy dissipation during PSII (Y[NPQ]), the photochemical quenching of PSII (qP), and the fraction of open PSII centers (qL) was calculated as follows according to Alexander V. Ruban (2017). However, Y(I), Y(ND), and Y(NA) were estimated according to Klughammer and Schreiber (1994). The electron transport rate (ETR) of PSII and PSI were calculated as ETR I (or ETR II) = 0.5 × abs I × Y(I) (or Y[II]), where 0.5 is the fraction of absorbed light between PSII and PSI (assuming they are equal), and abs I is absorbed irradiance taken as 0.84 of incident irradiance. All plant materials were kept in the dark for at least 30 minutes before being measured.

The determination of the CO_2_ assimilation rate was conducted with a portable gas-exchange system (LI-6400XT, Li-COR, Lincoln, NE). The photosynthetic rate was measured at a PPFD of 500 μmol photons m^−2^ s^−1^, a CO_2_ concentration of 400 μmol mol^−1^, relative humidity of 60%, and temperature of 25 °C.

### Quantifying photosynthetic pigments

To determine chlorophyll concentrations, six leaf disks were collected from each leaf sample with a leaf punch (φ = 5.35 mm) after the final harvest. They were then put into an Eppendorf tube containing 1 ml of 80% acetone solution. The chlorophyll was extracted without grinding after being incubated at 4 °C for 48 hours. The chlorophyll content was spectrophotometrically analyzed with a Spectrophotometer (Shimadzu, UV-1280) at wavelengths of 663 nm and 646 nm (Porra et al., 1989). To determine the above ground biomass, the above ground tissue of the plants was harvested, placed in paper envelopes, and dried at 80 °C for 48 hours. The dry weight was then determined with an analytical balance. SPAD values were determined with a hand-held SPAD-502 meter (Spectrum Technologies, Inc.). Anthocyanin concentrations were measured with a hand-held Anthocyanin Content Meter (CCM 200A plus, produced by V S INSTRUMENT PVT. LTD).

### Ascertaining the electron accepting abilities of the chemicals

Thylakoid membranes were prepared from spinach leaves to evaluate photosynthetic electron transport with the presence of chemicals (Terashima et al., 1989; Yamori et al., 2008). For preparation of thylakoid membranes, leaves were homogenized in an ice-cold blending buffer (50 mM Hepes-KOH, pH 7.8, 0.33 M sorbitol, 2 mM EDTA, 1 mM MgCl_2_, 1 mM MnCl_2_, 25 mM Na-ascorbate, and 1 mM DTT). The homogenate was filtered through a layer of Miracloth (Calbiochem) and the filtrate was centrifuged at 2,000 g for 30 seconds (s). The supernatant was removed and thylakoid membranes in the pellet were suspended with a suspending buffer (0.33 M sorbitol and 50 mM Hepes-KOH, pH 7.8). Electron transport activities in the thylakoid membranes (10 μg chlorophyll ml^−1^) suspended in the medium described above were measured with a Clark-type oxygen electrode (Hansatech, King’s Lynn, the UK) in a saturating white LED light, through monitoring the decreases or increases of O_2_ concentration with in the solution as explained by Yamasaki et al. (2002). To measure whole-chain electron transport activity, 5 mM methylamine, 0.5 mM sodium azide, and 50 μM methyl viologen (or 50 μM of chemicals) were added to the suspension. The assay medium for PSI electron transport activity contained 10 mM methylamine, 5 μM DCMU, 0.5 mM sodium azide, 150 μM DCIP, 1 mM sodium ascorbate, and 50 μM methyl viologen (or 50 μM of chemicals). Activity of PSII electron transport was determined with 50 μM of phenyl-p-benzoquinone (or 50 μM of chemicals) as an electron acceptor in the presence of 0.25 μM of 2,5-dibromo-3-methyl-6-isopropyl-p-benzoquinone and 1 μM of DBMIB, with modifications to the method of Yamasaki et al. (2002).

### RNA extraction and RNA-seq analysis

RNA was extracted from whole Arabidopsis plants after they were exposed to continuous light stress for 96 hours by using the RNeasy Plant kit (QIAGEN, Hilden, Germany). The library for RNA-seq was constructed using the protocol of Lasy-Seq (Kamitani et al., 2019) ver. 1.1 (https://sites.google.com/view/lasy-seq/) and sequenced using Nova Seq plus (Illumina, San Diego, CA, USA) with single-end sequencing lengths of 150 bp. All obtained raw reads were trimmed using the Trimmomatic ver. 0.33 (Bolger et al., 2014) with the following parameters: ILLUMINACLIP:TruSeq3-SE.fa:2:30:10 LEADING:20 TRAILING:20, SLIDINGWINDOW:4:15 MINLEN:40. The trimmed reads were mapped onto the reference A. thaliana genome (TAIR10; https://ftp.ensemblgenomes.ebi.ac.uk/pub/plants/release-55/fasta/arabidopsis_thaliana/dna/Arabidopsis_thaliana.TAIR10.dna.toplevel.fa.gz) using STAR ver. 2.7.10b (Dobin et al, 2013), and the read counts per genes were calculated using RSEM ver. 1.3.3 (Li and Deway,2011). Differentially expressed genes (DEGs) between the control group and the chemical treatment group were detected by the criteria of |fold-change| > 1.5 and false discovery rate = 0.1 using the DESeq2 R package ver. 1.40.2 (Love et al., 2014). Furthermore, the gene ontology (GO) enrichment analysis of DEGs was implemented by the criteria of adjusted P-value <0.01 using clusterProfiler R packages ver. 4.8.3 (Wu et al., 2021).

### Statistical analysis

The significance of variations between multiple means were evaluated by the Tukey–Kramer test. The significance of variations between two means were evaluated by the Student’s *t*-test. Statistical analysis was performed using Prism v. 8.0.1 software.

### Supplementary Data files

Two files are provided as supplementary materials:

*Supply 20240419* contains Supplementary Figure 1-7, and Supplementary Tabel 1-2.

*Supply table 20240419* contains Supplementary Table 1-2 in two sheets of excel format.

## Author Contributions and Acknowledgments

W.Y. conceived and designed the experiments. Y.Q. and K.S. mainly performed the experiments. Y.Q., K.S., M.N., T.A., I.T. and W.Y. analyzed the data, and Y.W. performed gene expression analysis. Y.Q. and W.Y. prepared figures and the manuscript. All authors have read and approved the final version of this manuscript.

This work was supported by KAKENHI to W.Y. (Grant Number: 18KK0170, 20K21346, 20H05687, 21H02171 and 24H02277) from Japan Society for the Promotion of Science (JSPS).

